# Touch-induced Mechanical Strain in Somatosensory Neurons is Independent of Extracellular Matrix Mutations in *C. elegans*

**DOI:** 10.1101/2019.12.27.889766

**Authors:** Adam L. Nekimken, Beth L. Pruitt, Miriam B. Goodman

**Affiliations:** Stanford University Department of Mechanical Engineering; Stanford University Department of Molecular and Cellular Physiology

## Abstract

Cutaneous mechanosensory neurons are activated by mechanical loads applied to the skin, and these stimuli are proposed to generate mechanical strain within sensory neurons. Using a microfluidic device to deliver controlled stimuli to intact animals and large, immobile, and fluorescent protein-tagged mitochondria as fiducial markers in the touch receptor neurons (TRNs), we visualized and measured touch-induced mechanical strain in *C. elegans* worms. At steady-state, touch stimuli sufficient to activate TRNs induce an average strain of 3.1% at the center of the actuator and this strain decays to near zero at the edges of the actuator. We also measured strain in animals carrying mutations affecting links between the extracellular matrix (ECM) and the TRNs but could not detect any differences in touch-induced mechanical strain between wild-type and mutant animals. Collectively, these results demonstrate that touching the skin induces local mechanical strain in intact animals and suggest that a fully intact ECM is not essential for transmitting mechanical strain from the skin to cutaneous mechanosensory neurons.

## Introduction

Touch and proprioception are essential to the daily lives of all animals, including humans. Classical examples of mechanotransduction, both of these senses depend on activation of mechano-electrical transduction (MeT) channels arrayed within somatosensory neurons (Katta et al., 2015) and are thought to activate following physical deformation of sensory cells during touch and movement. Consistent with this inference, neurons innervating stretch receptor organs in vertebrates (Bewick and Banks, 2015) and arthropods (Suslak and Jarman, 2015) are activated by experimentally applied mechanical strain (stretch). High-speed volumetric imaging reveals that proprioceptors deform in crawling Drosophila larvae (He et al., 2019; Vaadia et al., 2019) and that physical deformations are correlated with calcium transients. Similarly, movement induces localized calcium transients in the tertiary dendrites of the *C. elegans* PVD proprioceptors that depend on expression of a DEG/ENaC/ASIC protein (Tao et al., 2019). These studies reinforce the idea that physical deformation of sensory neurons during movement is critical for activation of native MeT channels, but do not address whether or not similar deformations occur in response to touch or measure the extent of mechanical strain that occurs within the sensory neurons themselves.

Using *C. elegans* touch receptor neurons (TRNs), we investigated the quantitative relationship between touch and physical deformation of sensory neurons in intact living animals. The ready availability of transgenic animals expressing TRN-specific markers and their transparent body make *C. elegans* an especially useful animal for investigating touch-evoked sensory neuron deformation. Adult animals have six TRNs, consisting of two bilaterally symmetric pairs of touch receptor neurons (ALM and PLM) and two neurons that run along the ventral midline (AVM and PVM) (Goodman, 2006). These six neurons extend long, unusually straight sensory neurites (Krieg et al., 2017) that are embedded in epidermal cells and have a distinctive, electron-dense extracellular matrix or ECM as well as hemidesmosome structures attaching the TRNs to the cuticle (Chalfie and Sulston, 1981; Chalfie and Thomson, 1979).

All of the TRNs express the MEC-4 channel, which is required for touch-evoked calcium transients (Suzuki et al., 2003) and touch-evoked MeT currents (O’Hagan et al., 2005). The MEC-4 channels localize to puncta arrayed along the entire length of wild-type TRN sensory neurites (Chelur et al., 2002; Cueva et al., 2007; Emtage et al., 2004; Katta et al., 2019), but are disrupted in ECM mutants (Emtage et al., 2004).

Touch-evoked behavior (Petzold et al., 2013) and MeT channel activation (Eastwood et al., 2015) depend on body indentation rather than the force applied. Slow stimuli fail to activate MeT currents and their size increases with stimulus frequency, indicating that activation of MeT channels in their native environment depends on tissue viscoelasticity (Eastwood et al., 2015; Katta et al., 2019; Sanzeni et al., 2019). Although slow, movement-induced physical deformations of the worm’s body and its neurons are too slow to activate MeT currents in the TRNs (Eastwood et al., 2015; Katta et al., 2019), these undulatory movements produce mechanical strains of up to 40% (Krieg et al., 2017; Krieg et al., 2014), indicating that TRNs can withstand significant mesoscale extension and compression *in situ*. The ability to withstand movement-induced strain is shared by mammalian nerves that experience up to 30% strain during limb movement (Phillips et al., 2004).

Here, we sought to determine whether or not body indentations sufficient to evoke calcium transients in TRNs also generate local strain. To achieve this goal, we visualized steady-state, touch-induced strain in TRNs in living animals restrained within a microfluidic stimulation device equipped with the ability to deliver mechanical stimuli (Fehlauer et al., 2018; Nekimken et al., 2017a). Our approach benefits from large, immobile mitochondria distributed within TRN sensory neurites (Sure et al., 2018), exploits the ability to tag these mitochondria with a red fluorescent protein (Zheng et al., 2014), and borrows analytic principles from traction force microscopy (Ribeiro et al., 2016). Although mitochondria have been used to evaluate neuronal mechanics in culture (O’Toole et al., 2015), we believe this study is the first to make use of mitochondria as mechanical fiducial markers in living animals. We show that body indentation increases local steady-state mechanical strain in the TRNs in a manner that is robust to mutations known to affect attachment of the TRN to the extracellular matrix and to epidermal cells.

## Results

### Mechanical stimulation in a microfluidic device

To directly observe touch-induced deformation of the TRNs, we sought to develop a method that combines the delivery of controlled mechanical stimuli and three-dimensional optical imaging. To reach this goal, we applied mechanical stimuli to adult worms confined in a microfluidic device (Nekimken *et al.*, 2017, Figure 1A). Our device has pneumatic actuators that consist of a thin flexible wall that separates a channel filled with air from a worm in the trap channel. When air pressure in the actuator channel is increased using a pressure controller, the thin wall expands like a balloon, deforming the trapped worm and generating indentations sufficient to activate the TRNs (Nekimken et al., 2017a). Other devices that deliver mechanical stimuli to restrained (Cho et al., 2018) or moving (McClanahan et al., 2017) worms in microfluidic chambers have been reported, but they use larger actuator channels and are thus not well-matched to our goal of investigating local deformation of the TRNs.

**Figure 1:**
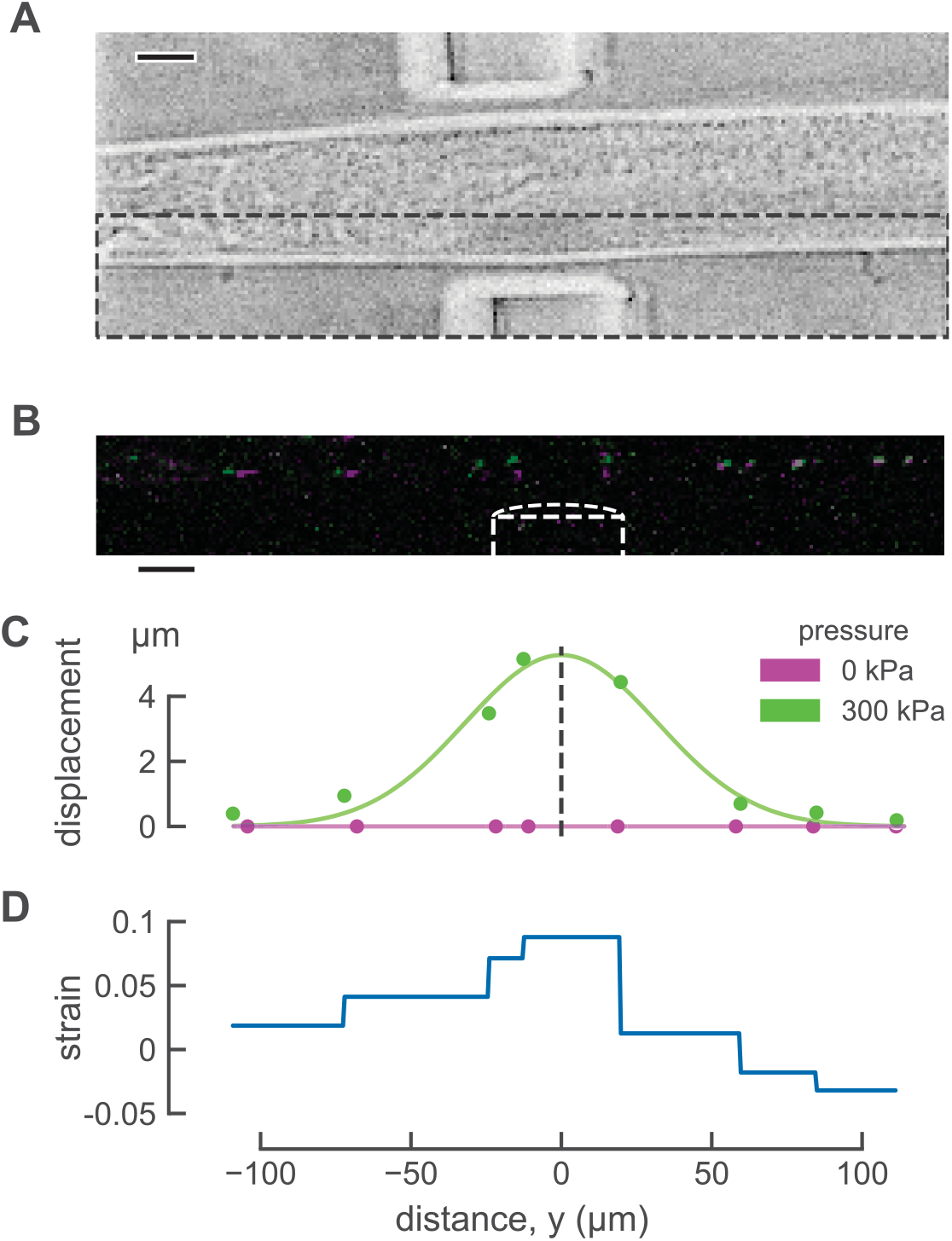
Touch-induced mechanical strain measured using mitochondria as fiducial markers in *C. elegans* TRNs *in vivo*. A) Brightfield image of worm in the microfluidic device. Scale bar 20 μm. B) Maximum projection image of TRN mitochondria before (magenta) and during (green) mechanical stimulus. Mitochondria that did not move appreciably appear white due to the overlap of magenta and green. Image intensity and contrast was adjusted to improve visualization. Scale bar 20 μm. C) Displacement in the direction of actuation before (magenta) and during (green) stimulation. The smooth line (green) is a Gaussian fit used to infer the center of actuator. D) Distribution of touch-induced strain in the TRN for a single actuation trial.

Our device is designed to increase the probability that animals will enter the trap with either their left or right sides in contact with the bottom of the chamber and their ventral and dorsal sides near the actuator (Nekimken et al., 2017a). Consistent with this expectation and the position of the AVM and ALM neurons within the worm’s body, previous experiments using this device resulted in a higher frequency of AVM activation compared to ALM (Nekimken et al., 2017a). However, the rotational orientation of each worm was variable. We took advantage of this variation to select animals oriented such that ALM (rather than AVM) was near the actuator and used only ALM neurons for this study. In all cases, this variation in worm positioning leads to variation in the size of the effective stimulus. To ensure that all animals received a deforming stimulus, we limited our analysis to animals whose TRNs were deformed enough to observe visually during the experiment.

### Mitochondria as fiducial markers for inferring strain

We used fluorescent protein-tagged mitochondria as fiducial markers to measure strain, an approach that uses particle tracking with large (~1μm), immobile mitochondria in the TRNs (Sure et al., 2018) serving as natural fiducial markers. We imaged mitochondria position before and after mechanical stimulation (Figures 1B and 1C). Next, we used the displacement between adjacent mitochondria to quantify one-dimensional mechanical strain along the long axis of the TRN (Figure 1D) according to: ɛ = Δ*L*/*L*_0_, where *L*_0_ is the resting, undeformed length of an object, and Δ*L* is the change in length of the object when deformed. For clarity, we refer to this as longitudinal strain. As shown in Figure 1, mitochondria adjacent to the actuator moved more than those anterior and posterior to the actuator (Figures 1B and 1C) and the inferred strain was greatest near the center of the actuator (Figure 1D). This method enables direct observation of touch-evoked strain in *C. elegans* TRNs in a single dimension aligned with the TRN’s longest dimension.

We note that this method is not suitable for detecting shear strain or bending strain. To derive these measurements, we would need to detect angular changes (shear) or relative position (bending) of the mitochondria within the TRNs. Given that the diameter of each mitochondrion is only ~100 nm and the TRN is diameter is not much larger than this (200-300 nm, see Figure 3C, Cueva et al., 2007), such movements cannot be resolved because these objects are similar in size to the estimated lateral resolution of 254 nm and axial resolution of 632 nm of our imaging system. What about strain in directions orthogonal to the long axis of the TRN dendrite? Although our image stacks contain three-dimensional position data (Figures 2A and 2B), the initial distance between adjacent pairs of mitochondria in the *x* and *z* directions is small compared to the resulting displacement. Thus, measurement resolutions lead to large uncertainties in the strain calculation for these dimensions. Both limitations can be attributed to the confinement of the mitochondria within the narrow caliber of the TRN and the fact that the TRNs lie primarily within a single focal plane. Thus, the size of mitochondria, the geometry of the TRNs and the optical resolution of the spinning-disk confocal limit this measurement to one-dimensional strain along the *y-*dimension that traces the main axis of the TRN.

**Figure 2:**
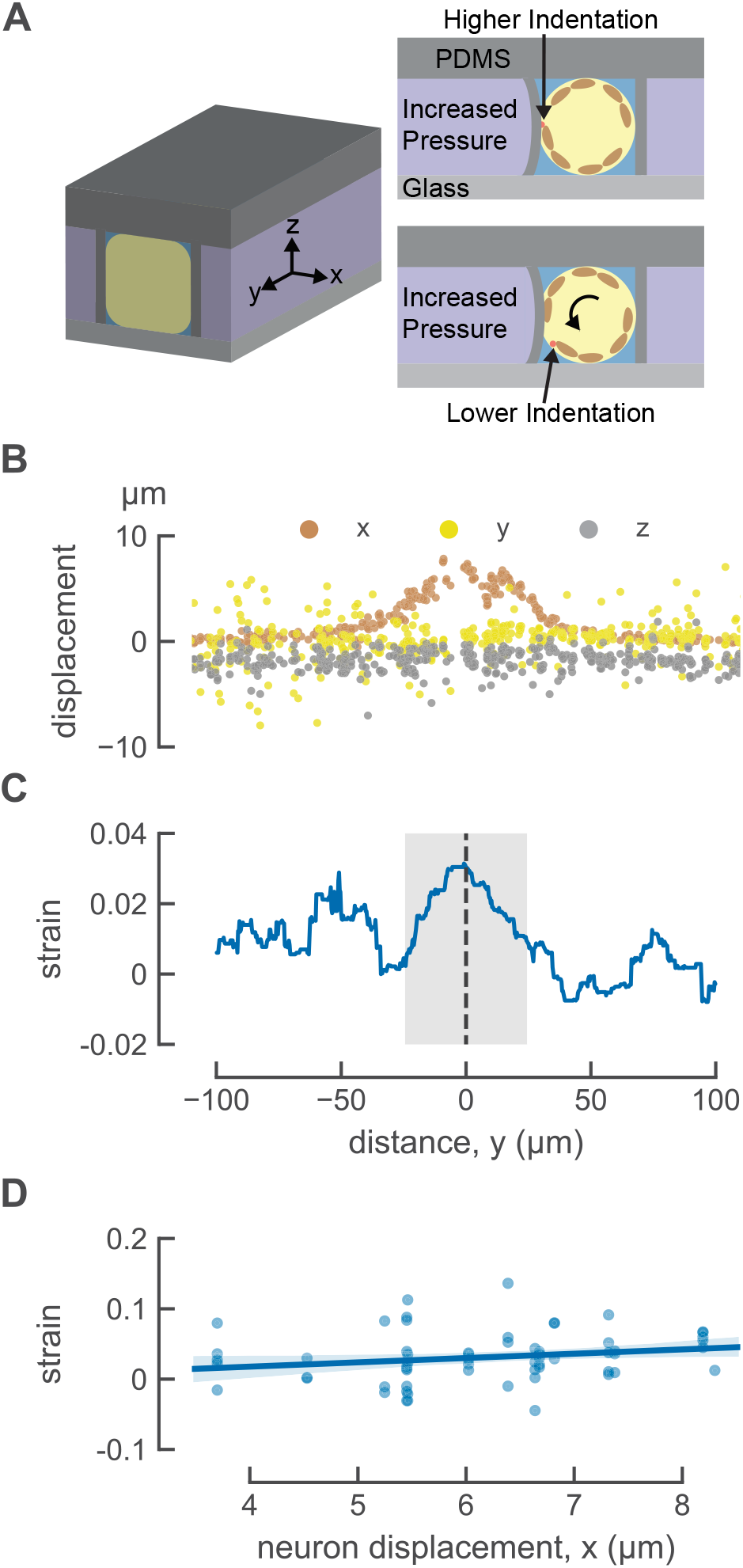
Indentation induces local longitudinal strain in *C. elegans* TRNs. A) Three-dimensional diagram the positioning of the worm in the microfluidics trap, the animal-centric coordinate system used to characterize strain, and the consequences of trapping animals in different orientations. TRN is red, and muscles are included in brown as visual aid. B) Touch-induced displacement in three dimensions. C) Touch-induced strain, anterior is to the left and posterior is to the right. *y=0* is the center of the actuator. *n*=61 trials from *N=*15 worms. D) Mechanical strain at *y=0* as a function of maximum neuron displacement. The line is a linear fit to the data: ε =0.0061x − 0.0067, where ε is the strain and x is the maximum displacement. The shaded area indicates 95% confidence intervals of the fit. *R*^2^ = 0.04.

**Figure 3:**
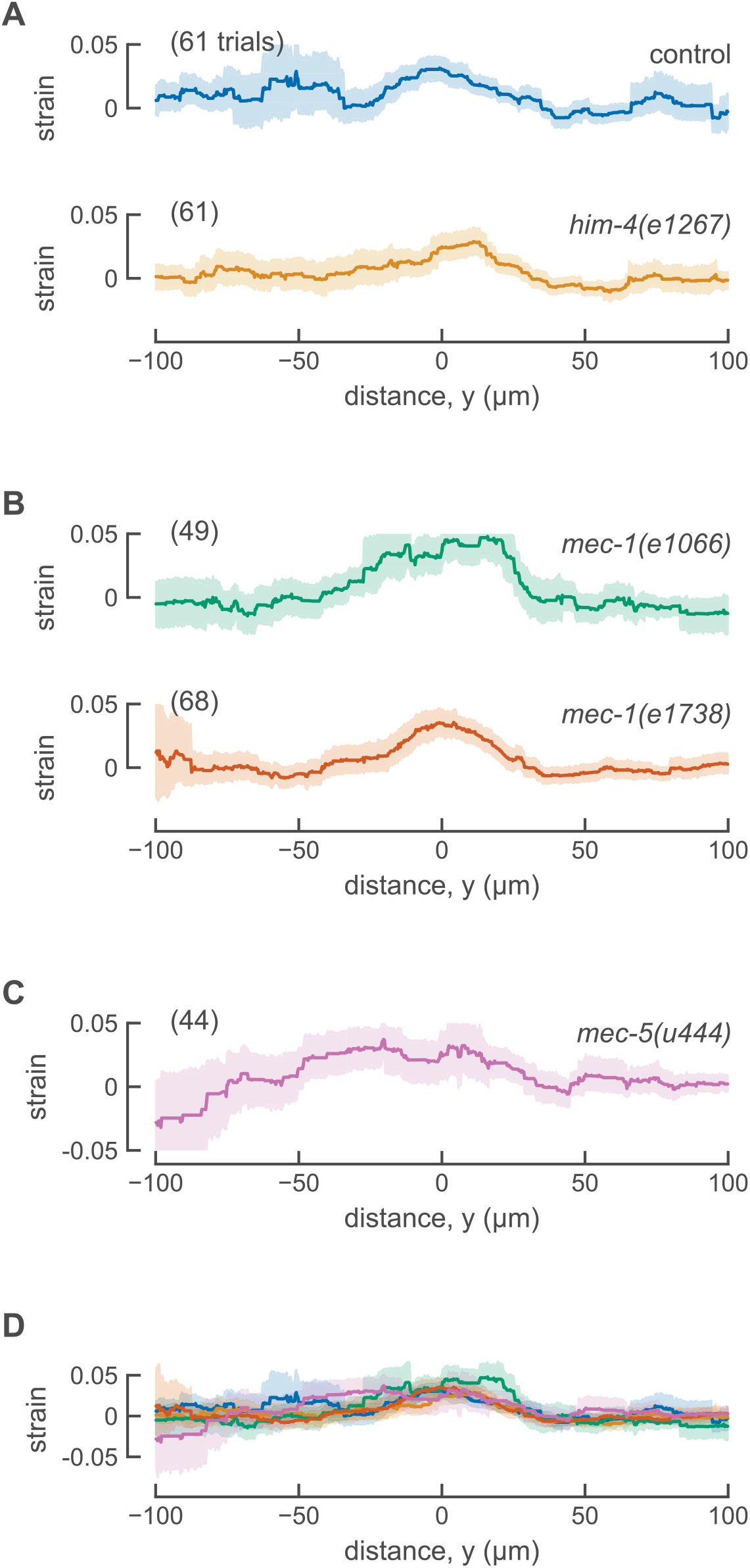
Touch-induced mechanical strain profiles are similar in control animals and ECM mutants. A) Spatially averaged strain in control ALM neurons with normal TRN attachment (15 animals) and *him-4(e1267)* with attachment defects (16 animals). The data for control animals are the same data as Figure 2. B) Spatially averaged strain in *mec-1* mutants. 14 independent animals for *mec-1(1066)* and 16 animals for *mec-1(e1738)*. C) Spatially averaged strain in *mec-5(u444)* mutants (17 animals). D) Overlay of all spatially averaged strain profiles. Smooth lines are the averages across all trials and shaded areas show the error (95% confidence intervals). One ALM neuron tested in each animal.

### Local indentation induces local mechanical strain in TRNs

Next, we estimated the spatially averaged distribution of touch-induced strain in the TRN. The mitochondria are sparsely distributed in the TRN and strain is a pairwise measurement between adjacent mitochondria, so our strain measurements result in a series of discontinuous step functions for a single trial rather than a smooth curve (Figure 1D). Each segment of the plot indicates the average strain between two markers, so the maximum local strain on each interval may be larger than our measured strain. The average distance between mitochondria in control *jsIs1073* animals was 26 μm (min: 5 μm; max: 64 μm) and we were able to analyze an average of 8.9 intervals per trial (Table S1). To obtain averages across trials, we plotted deformation against position in the longitudinal or *y* direction, fit this profile with a Gaussian function and defined the position of the maximum as the center of the actuator and *y*=0. (see dotted line in Figure 1C). In this coordinate system, positions anterior to *y=0* are negative and positions posterior to *y=0* are positive. Finally, we determined the average strain at each longitudinal position along the neurite across actuation trials (see Figure 2C). Note that the average strain is both larger and noisier for anterior (negative) positions. This is likely to reflect the fact that the head is less constrained in the microfluidic channel than other parts of the body and suggests that displacement and strain on the anterior side of the actuator include both touch-and movement-induced mechanical strain on the neuron.

As expected, strain at the center of the actuator (*y=0*) increases with TRN deformation (Figure 2D), but the dependence on TRN deformation was weak. The observed, but modest variation in TRN deformation arises from two aspects of our method. First, because the actuator is fixed to the surrounding material of the device on all sides, its center is the location of maximum deformation. Second, although most animals are trapped with either their left or right side in contact with the coverslip at the bottom of the device, some are rotated along their anterior-posterior axis such that the position of the imaged TRN varies with respect to the actuator (Figure 2A). Thus, the average strain induced between the pair of mitochondria at the center of the actuator was 0.031±0.005 (mean ± SEM, *n* = 61 trials, *N*=15 animals) in transgenic *jsIs1073* TRN neurons. Although additional studies exploring a wider range of deformations will be needed to determine the nature relationship between TRN deformation and strain, to our knowledge, these are the first *in vivo* measurements of touch-induced longitudinal strain in touch receptor neurons, and they link local body indentation to local cellular deformation.

### Do tagged mitochondria affect TRN function?

We used classical touch assays to address this question (Chalfie et al., 2014; Nekimken et al., 2017b) by measuring touch sensitivity in wild-type (N2) and transgenic worms carrying the transgenic mitochondria marker, *jsIs1073*. In blinded assays of three independent cohorts of 25 animals for each genotype, wild-type and *jsIs1073* animals had touch response rates of 0.869 and 0.865, respectively. The difference between the means was 0.00667 [95% CI (−0.0347, 0.048)]. Thus, the *jsIs1073* transgene does not decrease touch sensitivity.

### Selected ECM mutants and their touch sensation and anatomical phenotypes

Having measured touch-induced mechanical strain in control TRNs, next we measured longitudinal strain in four existing ECM mutants: *him-4(e1267), mec-1(e1738)*, *mec-1(e1066),* and *mec-5(u440).* One of these mutants, *him-4(e1267),* has a partial defect in touch-sensitivity, and the others are touch-insensitive (Du et al., 1996; Emtage et al., 2004; Vogel and Hedgecock, 2001). We reproduced these previous results using a ten-touch assay (Table 1). Whereas *him-4* mutants retain MEC-4 puncta that are grossly wild-type, TRN neurites in *mec-1* and *mec-5* mutants lack prominent MEC-4 puncta (Emtage et al., 2004). In *him-4* and in *mec-1(e1738)* mutants analyzed here, the ALM and PLM neurons are displaced from their normal body position near the lateral midlines and are not properly embedded in the epidermis (Emtage et al., 2004; Vogel and Hedgecock, 2001). This effect is inferred to arise from a defective TRN-ECM attachment (Vogel and Hedgecock, 2001). Consistent with this idea, the electron-dense ECM is not detected in *mec-1* mutants (Chalfie and Sulston, 1981). Analyzing these four mutants enables us to evaluate the relationship, if any, between touch-induced longitudinal strain and three other phenotypes: behavioral responses to touch, the distribution of MEC-4 puncta, and TRN-ECM attachment.

**Table 1:**
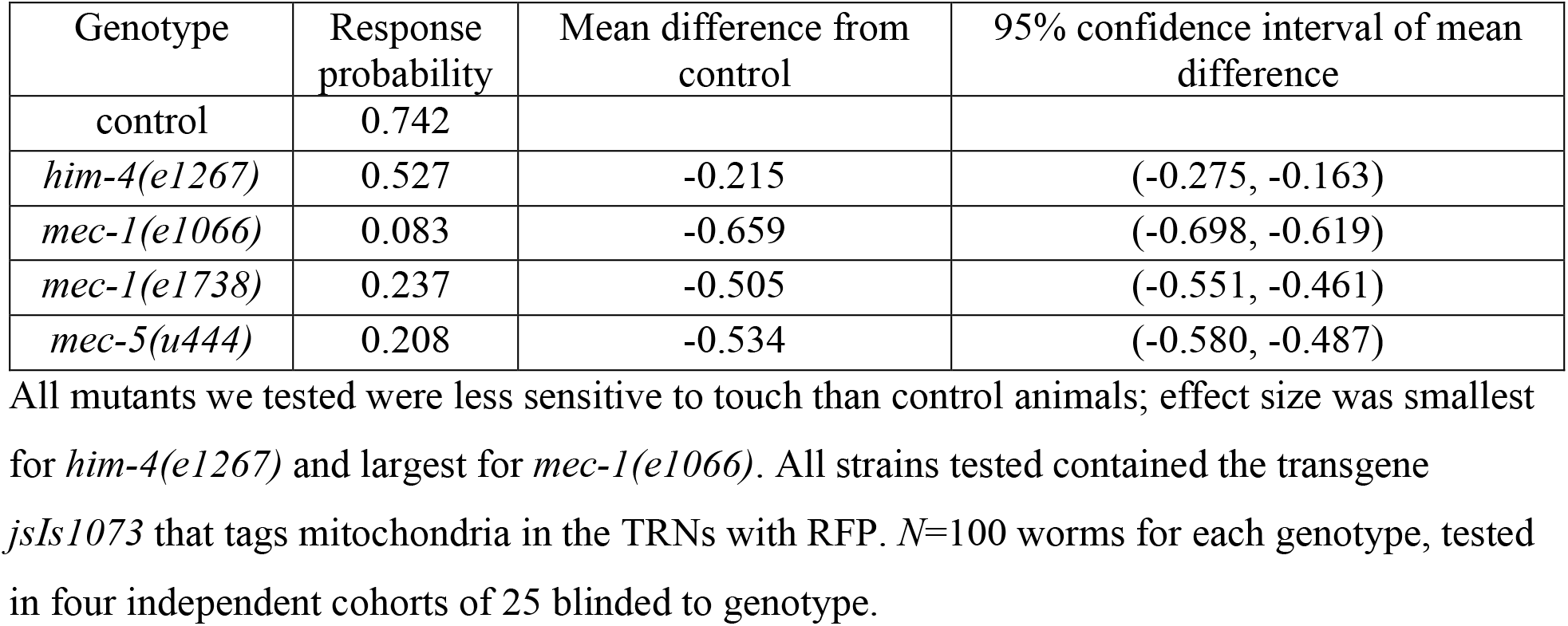
Touch sensitivity of ECM mutants.

The *him-4* gene encodes hemicentin, a conserved ECM protein that is rich in Ig and EGF domains and is expressed by body wall muscles (Vogel and Hedgecock, 2001). The *him-4* locus is >30 kb, which complicated determination of the molecular defect encoded by *e1267* using classical sequencing methods. We used whole-genome sequencing to circumvent this limitation and found that *e1267* encodes a single base indel in the third intron of the *him-4* gene, introducing a shift in the reading frame of the encoded protein (Figure S1). A second polymorphism was detected in the intron following the 48th exon. Thus, the *e1267* allele is a null allele and animals carrying this mutant are likely to lack the HIM-4 protein. The *mec-1* gene encodes a large secreted protein rich in kunitz-like domains, is expressed by the TRNs, and many alleles of this gene were found in forward genetic screens for touch-insensitive animals (Chalfie and Au, 1989; Chalfie and Sulston, 1981). The *e1066* allele is a null and animals carrying this allele are likely to lack the MEC-1 protein, whereas the *e1738* allele encodes a premature stop codon and is proposed to express a truncated MEC-1 protein. Both *mec-1* mutants are touch insensitive, but only *e1066* lacks proper TRN-ECM attachments (Emtage et al., 2004). *mec-5* encodes an atypical collagen and, unlike *mec-1*, it is not made by the TRNs (Du et al., 1996). Like other ECM mutants we analyzed here, the *u440* allele is null and *u440* mutants do not express the MEC-5 protein. The TRN neurites in *mec-5* mutants are positioned near the lateral midlines, but are less straight than they are in wild-type animals, meandering such that some segments of the neurite are close to the muscle and other parts closer to the lateral midline (Emtage et al., 2004).

### Touch-induced longitudinal strain in TRNs with ECM mutants

To analyze transmission of mechanical energy from the skin in animals lacking proper TRN-ECM attachment, we introduced the *jsIs1073* transgene into *him-4* and *mec-1* mutants and applied mechanical stimuli to animals trapped in our pneumatic microfluidic device. Touch-induced longitudinal mechanical strain in *him-4(e1267)* TRNs was indistinguishable from that observed in control animals (Figure 3A, Table 2). Touch-induced longitudinal strain was likewise similar to control in both *mec-1* mutants (Figure 3B) and in *mec-5* mutants (Figure 3C). Figure 3D shows that the average longitudinal strain profiles for control, *him-4*, *mec-1*, and *mec-5* mutants are similar across the entire 200 μm segment of the TRN neurite analyzed here. Values for strain measured at the actuator center (*y=*0) had similar mean values and distribution for all genotypes tested (Figure 4A) and estimation graphics (Ho et al., 2019) indicate that the mean values for all genotypes are not different than control values (Figure 4B). Except for *mec-5* mutants, the number of mitochondria available for tracking and the average distance between mitochondria was similar in control transgenic and mutant animals (Table S1). *mec-5* mutants had fewer, more widely spaced mitochondria than control animals, an effect would be expected to impair the spatial resolution of longitudinal strain measurements. Collectively, these findings suggest that neither severe nor mild defects in TRN-ECM attachment play a significant role in the generation of touch-induced longitudinal mechanical strain at steady-state. Future studies and techniques will be needed to resolve dynamic changes in local strain that are expected to occur on the millisecond timescale of MEC-4-dependent MeT channel activation.

**Table 2:** Touch-evoked longitudinal strain in the TRNs as a function of genotype.

| Genotype | Mean strain ± SEM | Mean difference from control | 95% confidence interval of mean difference | stimulation trials | animals |
| --- | --- | --- | --- | --- | --- |
| control | $0.031 \pm 0.005$ | | | 61 | 15 |
| *him-4(e1267)* | $0.024 \pm 0.005$ | -0.007 | (-0.020, 0.007) | 61 | 16 |
| *mec-1(e1066)* | $0.034 \pm 0.007$ | 0.003 | (-0.012 0.022) | 49 | 14 |
| *mec-1(e1738)* | $0.035 \pm 0.005$ | 0.004 | (-0.009 0.018) | 68 | 16 |
| *mec-5(u444)* | $0.026 \pm 0.009$ | -0.005 | (-0.028 0.013) | 44 | 17 |

There was no detectable difference in the magnitude of the touch-evoked strain in control and ECM mutant animals.

**Figure 4:**
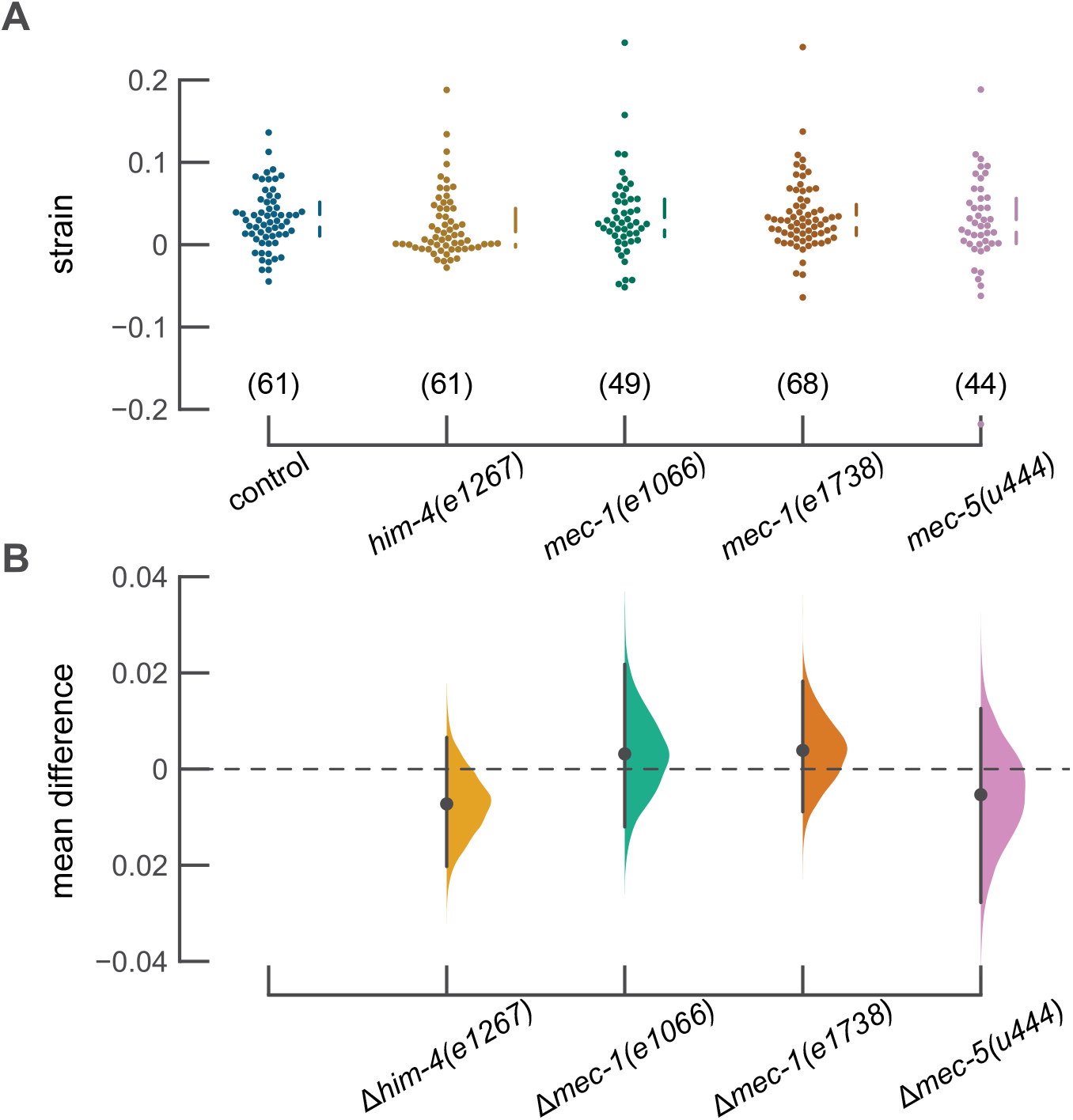
Strain at the center of the actuator is similar in control animals and ECM mutants. A) Strain at the center of the actuator. Each point is the result of a single trial. The vertical lines next to the swarm plots indicate the median and quartiles of the data. Data collected from 15, 16, 14, 16, and 17 animals (left to right). B) Difference in the mean strain between control and each mutant with a bootstrapped resampled distribution of the data. Estimation plots generated using the DABEST plotting package (Ho et al., 2019).

## Discussion

Using mitochondria as natural fiducial markers in *C. elegans* TRNs, we showed for the first time that body indentation sufficient to activate the TRN causes an increase in longitudinal strain. The magnitude of this strain increases with TRN displacement. In wild-type animals, the touch-induced strain closest to the point of maximum indentation was roughly 3% (Figure 3, Table 2). This change in strain is unlikely to damage these sensory neurons, since they are subjected to mechanical strain on the order of 40% during locomotion (Krieg et al., 2017; Krieg et al., 2014). Such global changes in strain are too slow to activate the TRNs (Eastwood et al., 2015; Katta et al., 2019).

The 3% strain we measured may be a lower bound on the true value for longitudinal strain. Simulations predicted an elongated strain field with a peak value of 12% for longitudinal strain (Sanzeni et al., 2019). Unlike our measurements, which used a wide (50μm), flexible PDMS membrane to indent the worm’s body and have limited spatial resolution, the simulations employed a stiff spherical bead (10μm diameter) to deliver touch stimuli and a continuous deformation function. The simulations (Sanzeni et al., 2019) predict strains are present in the other two dimensions (along the direction of the stimulus and tangential to the circumference of the worm at the point of the stimulus). Due to the nature of our measurement, however, we were only able to measure one-dimensional strain along the length of the TRN. Nevertheless, these measurements of touch-induced strain can be incorporated into future models of *C. elegans* touch sensation to improve understanding of the mechanical state of the TRN and mechano-sensitive ion channel complex upon touch stimulation.

### Origins of mechanical coupling between the skin surface and somatosensory neurons

We tested the idea that strain transmission would depend on TRN-ECM attachment, be correlated with impaired touch sensation, and be independent of the proper distribution of MEC-4 channels. To our surprise, we found that touch-induced longitudinal mechanical strain was similar in control TRNs and all ECM mutants tested here, including the *him-4(e1267)* and *mec-1(e1066)* mutants exhibiting severe defects in TRN-ECM attachments. Thus, touch-induced longitudinal strain we observed in the ALM neurons is independent of their attachment to other tissues or the expression of ECM proteins MEC-1 and MEC-5, at least at steady state.

This finding implies that mechanical coupling between the skin and sensory neurons persists in mutants with ECM defects. Friction is one alternative source of mechanical coupling between TRNs and surrounding tissues. In this scenario, all of the worm’s tissues are compacted together by a high hydrostatic pressure and this would elevate friction between tissues. Further compression applied to the outside of the worm during touch stimulation could lead to stiffening of the worm’s body that might be caused by internal structures jamming together (Gilpin et al., 2015). When we and others immobilize worms in microfluidic devices, we fabricate channels small enough to apply gentle compressive forces that the worm cannot overcome. As a result, the worm is compressed on all sides except the nose and tail when in the microfluidic trap, potentially jamming the TRNs against surrounding tissues and further increasing friction. Another possibility is that the generation of touch-induced local mechanical strain is dominated by the cytoskeleton. In support of this idea, mutations that disrupt the expression of MEC-12 α-tubulin and MEC-7 β-tubulin decrease mechanoreceptor currents and increase stimulus amplitude needed to activate these currents (Bounoutas et al., 2009; O’Hagan, 2005). Additional experimental work and methods with improved spatial and temporal resolution will be needed to differentiate among these possibilities.

### Spatial and temporal resolution of mechanical strain transmission

The spatial resolution of our strain measurements is limited by the average distance between adjacent, immobile mitochondria, which was 26μm (Table S1). Thus, this method may lack sufficient spatial resolution to detect micro- or nanoscale variations in mechanical strain transmission at steady state. The spatial resolution of strain measurements could be improved by using more closely spaced fiducial markers. In principle, MEC-4 channels (with an average spacing of 2-3 μm) tagged with a very bright fluorescent protein would provide a ten-fold increase in resolution. Independent of potential improvements in spatial resolution, the present approach is limited to steady-state measurements because our volume imaging rate is low (15 seconds per stack). In this regard, it is important to note that wild-type *C. elegans* TRNs are preferentially activated mostly by high-velocity stimuli (Eastwood et al., 2015; Katta et al., 2019; Nekimken et al., 2017a; Suzuki et al., 2003). Temporal resolution could be improved using other imaging techniques enabling rapid acquisition of imaging volumes. If volumes could be acquired at 20 Hz or faster, it would possible to observe mechanical strain in a 10 Hz buzz stimulus that we previously used to activate the TRNs (Nekimken et al., 2017a).

### Conclusion

We performed the first *in vivo* measurements of touch-induced mechanical strain in *C. elegans* TRNs. We used mechanically stable mitochondria in the TRNs as natural fiducial markers to observe deformation of the TRN and found that local touch stimuli applied in a microfluidic device induces local strain in the TRN. Defects in the ECM surrounding the TRN did not alter the steady-state mechanical strain in the TRN, suggesting that explicit attachments are not necessary for deformation of the TRN and that the bulk properties of tissues are sufficient to sustain significant mechanical energy transfer from the skin surface to the embedded neurons. In light of the temporal limitations of our measurements, however, we cannot exclude the possibility that TRN-ECM attachments contribute to dynamic aspects of mechanical strain transmission. Collectively, these findings provide an empirical basis for the idea that mechanical stimuli applied to the skin stretch embedded sensory neurons that may be shared by the sensory neurons that innervate the skin and other tissue in mammals.

## Methods and Materials

### Nematode strains

For all experiments measuring mechanical strain, we used animals carrying *jsIs1073* [mec-7p∷TagRFP-mito∷CBunc-119], a transgene that drives expression of TagRFP in the mitochondria in the TRNs (Zheng et al., 2014), alone or together with ECM mutants (Table 3). We relied on visualization of the *jsIs1073* transgene and behavioral phenotypes (for *mec* genes) or anatomical defects (for *him-4*) to perform genetic crosses. We used gene sequencing to verify the presence of ECM mutants in all strains created for this study. The primer pairs we used to amplify the relevant segments of each gene are:

*mec-1(e1066)*: Forward–catcttccacgccgcaaagtc, Reverse—aatcctctctgccctcatgttcc
*mec-1(e1738)*: Forward—tcacagtcagacgtgcctcg, Reverse—cattgcctcacaccaacttccac
*mec-5(u444)*: Forward—cagaatactatgtacgtaacttgggatc; Reverse—ctcatgggtacgcaaatgatactc
*him-4(e1267)*: Forward—tttcgtgatgactggtgactgtgg; Reverse—ttaaagtcaacagcaccgtgacc

**Table 3:** *C. elegans* strains.

| Strain Name | Genotype | Source |
| --- | --- | --- |
| N2 (Bristol) | wild-type | CGC |
| NM3573 | *jsIs1073* | (Zheng et al., 2014) |
| GN885 | *jsIs1073;him-4(e1267)* | this study |
| GN886 | *jsIs1073;mec-1(e1066)* | this study |
| GN887 | *jsIs1073;mec-1(e1738)* | this study |
| GN906 | *jsIs1073;mec-5(u444)* | this study |

### Immobilization and mechanical stimulation with a pneumatic microfluidics device

To provide a repeatable mechanical stimulus that is compatible with imaging the mitochondria of the TRN, we used a microfluidic device made of the transparent elastomer PDMS for simultaneous mechanical stimulation and imaging of *C. elegans* (Nekimken et al., 2017a). Using a pressure controller (Elveflow OB1), we applied 300 kPa of pressure to one of the device’s actuators to create a mechanical stimulus. We fabricated and operated devices designed for use with young adult worms as described in our previous work (Fehlauer et al., 2018; Nekimken et al., 2017a). Because worms immobilized in this trap are uniformly and partially deformed by the channel, we refer to this as the rest configuration.

### Sample preparation and inclusion criteria

We performed all experiments using young adult animals that were synchronized by hypochlorite treatment (Stiernagle, 2006) and cultured for three days on NGM agar plates seeded with OP50 bacteria for food. We transferred animals from the agar plates with a platinum wire pick to a drop of imaging medium (see below) in a small Petri dish. After using the pick to push away large particles that might clog the microfluidic device, we aspirated worms into polyethylene tubing (PE50, Intramedic™ brand, Becton-Dickson) with a syringe and connected the tubing to the device’s inlet. For each trial, we pushed a worm into the trap channel using the syringe and then evaluated whether to use this worm for an experiment based on our inclusion criteria. We included animals that fit in the microfluidic trap with minimal movement and could be ejected through the narrow opening at the head of the trap. When performing experiments with *him-4* mutants, we chose animals whose gross morphology was wild-type and avoided animals whose intestines were everted. In all cases, we only acquired images from animals whose ALM neurons were oriented adjacent to the mechanical actuation channels.

To improve image quality by reducing reflections generated by the walls of the microfluidic trap, we designed a non-toxic imaging medium with a refractive index similar to PDMS (1.4). The imaging medium was a 70%:30% (vol:vol) mixture of physiological saline and iodixanol (Optiprep™, Sigma-Aldrich), a non-toxic density-gradient medium (Boothe et al., 2017). We used the same physiological saline as that used for electrophysiological recordings from *C. elegans* neurons (O’Hagan et al., 2005), which contains (in mM): NaCl (145), KCl (5), MgCl_2_ (5), CaCl_2_ (1), and Na-HEPES (10), adjusted to a pH of 7.2 with NaOH. The imaging medium has an osmolality of 300-325 mOsm (Fiske Micro-Osmometer Model 210), so it does not cause large osmotic shocks to *C. elegans*. By contrast with other fluids we tested (e.g. glycerol, halocarbon oil), this medium has a viscosity that appeared to be similar to that of physiological saline.

### Image acquisition

Although the mechanical stimulus is mostly in the horizontal direction, there is enough deformation in the z-direction to move the neuron out of plane during stimulation in some cases. In initial experiments with a traditional epifluorescence microscope, the mitochondria often moved out of focus during stimulation due to movement in the z-direction. To account for this problem, we used a spinning disk confocal microscope (Nikon TiE, Yokogawa CSU-X1, 40x/NA 1.4 oil objective, and Photometrics Prime95B sCMOS camera) to acquire z-stacks, which provided adequate resolution in the z-direction.

For each trial, we acquired eleven (11) z-stacks containing the neuron of interest, with 300 kPa of pressure applied during the even-numbered stacks, and 0 kPa applied during the odd-numbered stacks. We started acquisition of a stack every 15 seconds, although the time to acquire each given stack was approximately 12 seconds. The exact time varied depending on the height of the z-stack, which was manually set to accommodate observed motion in the z direction during the test actuation. During the short delay between acquisition of stacks, we toggled the applied pressure.

### Image analysis

To detect the mitochondria, we used a particle-tracking algorithm implemented in Python (Allan et al., 2018), based on work by Crocker and Grier (Crocker and Grier, 1996). Briefly, the algorithm involves applying a spatial band pass filter, finding peaks, refining the position of peaks by finding their center of mass, and linking particles across timepoints into trajectories. In some stacks, the algorithm failed to detect the mitochondria because they were too close to the top or bottom of the stack, were not bright enough, overlapped with the mitochondria of another TRN, or were blurred due to the motion of the worm. We discarded all stacks subsequent to a stack where the image processing failed, because the strain measurement requires comparison to a previous stack. Additionally, not all of the stacks from each trial were usable, since the fluorophores bleached over time. As a result, not all actuation events yielded strain measurements (see Supplementary Figure S2).

Due to variability across trials, we manually selected a region of interest around the TRN and tuned parameters in the particle tracking algorithm. Primarily, we changed the minmass threshold, which filters out particles where the sum of pixel values within the boundaries of the particle is below the chosen threshold, and the search radius for the linking step, which specifies how far a particle can travel between timepoints and still be identified as the same particle. Less frequently, we changed the cutoff size for the bandpass filter or the brightness percentile threshold, which sets a minimum value for the brightest pixel in a particle as a percentile of the brightness of pixels in the image. We tuned these parameters until the particles found and linked included only particles along the location of the neuron and not autofluorescent spots. Supplemental Figure S3 shows the range of parameters we used.

### Touch assays

To test the touch-sensitivity of the mutants used in our strain transmission experiments, we performed touch assays by lightly stroking an eyebrow hair across the body of a worm and scoring its behavioral response (Goodman, 2006). For each session of touch assays, we tested 25 worms from a plate, performing 10 touches per worm. We performed the touch assays blinded with respect to genotype. For each touch event, we counted a response consisting of reversing direction or speeding up to move away from the stimulus as a positive response.

### *Whole-genome sequencing of* him-4(e1267)

Our whole-genome sequencing protocol involved four sub-protocols: 1) DNA extraction, 2) sequencing library preparation, 3) sequencing, and 4) analysis. We isolated DNA from CB1267 *him-4(e1267) X* animals with a phenol chloroform isoamyl alcohol (PCI) extraction. First, we washed worms off a mostly starved plate using M9 buffer, rinsed the animals twice in M9 buffer, resuspended them in EN buffer (0.1M NaCl and 20 mM EDTA), removed the supernatant, and flash-froze the sample in liquid nitrogen. Next, we added 450 μL of worm lysis buffer (0.1 M TRIS pH 8.5, 0.1 M NaCl, 50 mM EDTA, and 1% SDS) and 40 μL of proteinase K (10 mg/ml) to 50 μL of frozen worms and incubated at 62°C for 45 minutes, vortexing occasionally. Then, we performed the PCI extraction in a phase lock gel tube (VWR) by adding 500 μL of PCI to the sample, vortexing, spinning for 5 minutes at 10,000 rpm, and then collecting the upper phase. We repeated the PCI extraction step, and then extracted twice using chloroform. We precipitated the DNA by adding 40 μL of 5M sodium acetate and 1 mL of ethanol, spinning for 5 minutes at 10,000 rpm, removing the supernatant, washing in 70% ethanol, and resuspending in 50 μL of TE buffer at pH 7.4.

We created a sequencing library according to manufacturer instructions (Nextera DNA Library Prep Kit, Illumina). Briefly, this includes tagmentation of the DNA using the Illumina Tagment DNA Buffer and Enzyme, clean-up of the tagmented DNA using a Zymo DNA Clean and Concentrator Kit, PCR to add the library indices to the tagmented DNA, and PCR cleanup by gel extraction using a Qiagen MinElute Kit. We did a quality control step to confirm that the average length of DNA fragments was in the expected range of 300-500 bp using the Agilent Bioanalyzer at the Stanford Protein and Nucleic Acid Biotechnology Facility. Sequencing was completed on an Illumina NextSeq sequencer in the Stanford Functional Genomics Facility.

We analyzed the sequencing data using the computing cluster of the Stanford Center for Genomics and Personalized Medicine. Briefly, we mapped reads using Bowtie2, used Picard to sort reads, mark duplicates, and prepare read groups, then used GATK to select high quality SNPs and INDELs, and SnpEff to annotate the results.

## Abbreviations

TRN: touch receptor neuron
ECM: extracellular matrix

## Code availability

Two kinds of data were analyzed using custom code: image analysis (Python, https://github.com/anekimken/SSN_ImageAnalysis) and whole-genome sequence analysis available (scripts, https://github.com/wormsenseLab/whole_genome_sequencing).

## Acknowledgments

We thank Chloe Girard, Chantal Akerib, Melissa Pickett, and Anne Villenueve for the gift of their training and assistance in whole genome sequencing; Michael Nonet for NM3573, the strain with labeled mitochondria in the TRNs; Ehsan Rezeai for help with software testing; Michael Lin for the loan of a spinning disk confocal for pilot experiments. Some of the experiments were performed at Stanford core facilities, including the Cell Sciences Imaging Facility, Functional Genomics Facility (funded in part by a Shared Instrumentation Grant, NIH S10OD018220), the Genetics Bioinformatics Service Center, and the nano@Stanford labs (supported by the National Science Foundation as part of the National Nanotechnology Coordinated Infrastructure under award ECCS-1542152). This work was funded by an NIH fellowship to ALN (F31NS100318) and grant to MBG (R35NS105092).

---

Figure S1: *him-4(e1267)* is likely to be a null allele. A) Map of *him-4* with *e1267* allele annotated. B) Close-up of insertion in sequence as indicated by dotted lines in Panel A. C) Sequences of both Indels found by sequencing. The first indel is likely to cause the null phenotype because it causes a frameshift in an early exon, whereas the second indel is in a later intron.

Figure S2: Quality control for image (z) stacks as a function of genotype. Eleven (11) image stacks were collected from each TRN analyzed for touch-induced strain. This plot shows the number of stacks that passed quality for each experiment. Note: The final stack was not used as it was acquired in the resting configuration and could not be compared to a subsequent stack collected in the indented configuration.

Figure S3: We tuned parameters of the particle tracking algorithm to account for variability across images.

**Table S1:**
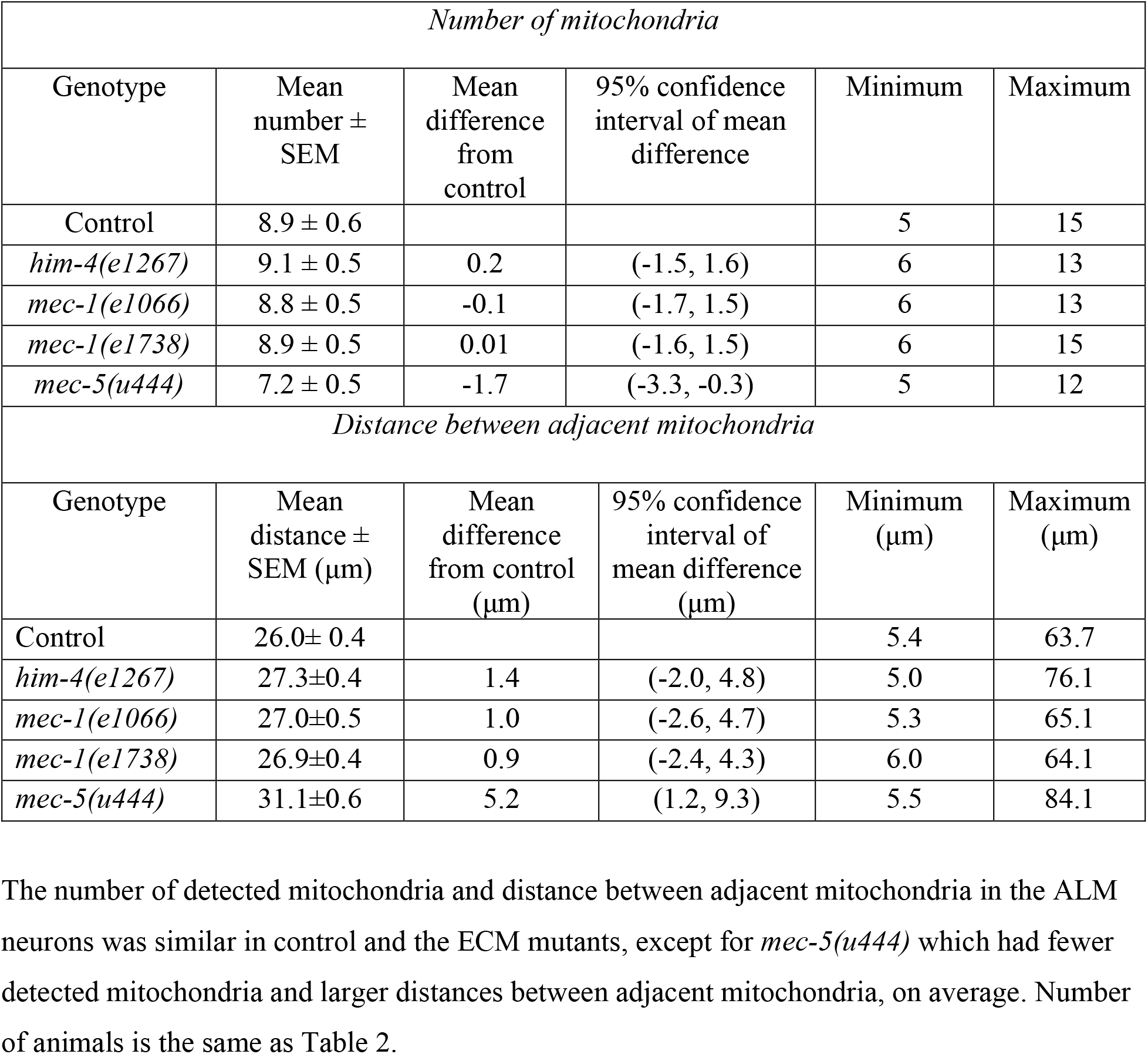
Average number of detected mitochondria and distance between adjacent detected mitochondria as a function of genotype.

| *Number of mitochondria* |  |  |  |  |  |
| --- | --- | --- | --- | --- | --- |
| Genotype | Mean number ± SEM | Mean difference from control | 95% confidence interval of mean difference | Minimum | Maximum |
| Control | 8.9 ± 0.6 | | | 5 | 15 |
| *him-4(e1267)* | 9.1 ± 0.5 | 0.2 | (-1.5, 1.6) | 6 | 13 |
| *mec-1(e1066)* | 8.8 ± 0.5 | -0.1 | (-1.7, 1.5) | 6 | 13 |
| *mec-1(e1738)* | 8.9 ± 0.5 | 0.01 | (-1.6, 1.5) | 6 | 15 |
| *mec-5(u444)* | 7.2 ± 0.5 | -1.7 | (-3.3, -0.3) | 5 | 12 |
| *Distance between adjacent mitochondria* |  |  |  |  |  |
| Genotype | Mean distance ± SEM (μm) | Mean difference from control (μm) | 95% confidence interval of mean difference (μm) | Minimum (μm) | Maximum (μm) |
| Control | 26.0± 0.4 | | | 5.4 | 63.7 |
| *him-4(e1267)* | 27.3±0.4 | 1.4 | (-2.0, 4.8) | 5.0 | 76.1 |
| *mec-1(e1066)* | 27.0±0.5 | 1.0 | (-2.6, 4.7) | 5.3 | 65.1 |
| *mec-1(e1738)* | 26.9±0.4 | 0.9 | (-2.4, 4.3) | 6.0 | 64.1 |
| *mec-5(u444)* | 31.1±0.6 | 5.2 | (1.2, 9.3) | 5.5 | 84.1 |

## References

Allan, D.B., Caswell, T., Keim, N.C., van der Wel, C.M., 2018. trackpy: Trackpy v0.4.1. doi:10.5281/ZENODO.1226458

Bewick, G.S., Banks, R.W., 2015. Mechanotransduction in the muscle spindle. Pflügers Arch. - Eur. J. Physiol. 467, 175–190.

Boothe, T., Hilbert, L., Heide, M., Berninger, L., Huttner, W.B., Zaburdaev, V., Vastenhouw, N.L., Myers, E.W., Drechsel, D.N., Rink, J.C., 2017. A tunable refractive index matching medium for live imaging cells, tissues and model organisms. Elife 6. doi:10.7554/eLife.27240

Bounoutas, A., O’Hagan, R., Chalfie, M., 2009. The Multipurpose 15-Protofilament Microtubules in C. elegans Have Specific Roles in Mechanosensation, Current Biology. doi:10.1016/j.cub.2009.06.036

Chalfie, M., Au, M., 1989. Genetic control of differentiation of the Caenorhabditis elegans touch receptor neurons. Science (80-.). 243, 1027–1033.

Chalfie, M., Hart, A.C., Rankin, C.H., Goodman, M.B., 2014. Assaying mechanosensation. WormBook.

Chalfie, M., Sulston, J., 1981. Developmental genetics of the mechanosensory neurons of Caenorhabditis elegans. Dev. Biol. 82, 358–370. doi:10.1016/0012-1606(81)90459-0

Chalfie, M., Thomson, J.N., 1979. Organization of neuronal microtubules in the nematode Caenorhabditis elegans. J. Cell Biol. 82, 278–289.

Chelur, D.S., Ernstrom, G.G., Goodman, M.B., Yao, C.A., Chen, L., O Hagan, R., Chalfie, M., O’Hagan, R., Chalfie, M., 2002. The mechanosensory protein MEC-6 is a subunit of the C. elegans touch-cell degenerin channel. Nature 420, 669–673. doi:10.1038/nature01205

Cho, Y., Oakland, D.N., Lee, S.A., Schafer, W.R., Lu, H., 2018. On-chip functional neuroimaging with mechanical stimulation in: Caenorhabditis elegans larvae for studying development and neural circuits. Lab Chip 18, 601–609. doi:10.1039/c7lc01201b

Crocker, J.C., Grier, D.G., 1996. Methods of Digital Video Microscopy for Colloidal Studies. J. Colloid Interface Sci. 179, 298–310. doi:10.1006/jcis.1996.0217

Cueva, J.G., Mulholland, A., Goodman, M.B., 2007. Nanoscale Organization of the MEC-4 DEG/ENaC Sensory Mechanotransduction Channel in Caenorhabditis elegans Touch Receptor Neurons. J. Neurosci. 27, 14089–14098.

Du, H., Gu, G., William, C.M., Chalfie, M., 1996. Extracellular proteins needed for C. elegans mechanosensation. Neuron 16, 183–194.

Eastwood, A.L., Sanzeni, A., Petzold, B.C., Park, S.-J., Vergassola, M., Pruitt, B.L., Goodman, M.B., 2015. Tissue mechanics govern the rapidly adapting and symmetrical response to touch. Proc. Natl. Acad. Sci. 112, E6955–E6963. doi:10.1073/pnas.1514138112

Emtage, L., Gu, G., Hartwieg, E., Chalfie, M., 2004. Extracellular proteins organize the mechanosensory channel complex in C. elegans touch receptor neurons. Neuron 44, 795–807.

Fehlauer, H., Nekimken, A.L.A.L., Kim, A.A.A.A., Pruitt, B.L.B.L., Goodman, M.B.M.B., Krieg, M., 2018. Using a microfluidics device for mechanical stimulation and high resolution imaging of C. Elegans. J. Vis. Exp. 2018. doi:10.3791/56530

Gilpin, W., Uppaluri, S., Brangwynne, C.P., 2015. Worms under Pressure: Bulk Mechanical Properties of C. elegans Are Independent of the Cuticle. Biophys. J. 108, 1887–1898.

Goodman, M., 2006. Mechanosensation. WormBook. doi:10.1895/wormbook.1.62.1

He, L., Gulyanon, S., Mihovilovic Skanata, M., Karagyozov, D., Heckscher, E.S., Krieg, M., Tsechpenakis, G., Gershow, M., Tracey, W.D., 2019. Direction Selectivity in Drosophila Proprioceptors Requires the Mechanosensory Channel Tmc. Curr. Biol. 29, 945–956.e3. doi:10.1016/j.cub.2019.02.025

Ho, J., Tumkaya, T., Aryal, S., Choi, H., Claridge-Chang, A., 2019. Moving beyond P values: data analysis with estimation graphics. Nat. Methods. doi:10.1038/s41592-019-0470-3

Katta, S., Krieg, M., Goodman, M.B., 2015. Feeling Force: Physical and Physiological Principles Enabling Sensory Mechanotransduction, in: Annual Review of Cell and Developmental Biology. doi:10.1146/annurev-cellbio-100913-013426

Katta, S., Sanzeni, A., Das, A., Vergassola, M., Goodman, M.B., 2019. Progressive recruitment of distal MEC-4 channels determines touch response strength in C. elegans. J. Gen. Physiol. doi:10.1085/jgp.201912374

Krieg, M., Dunn, A.R., Goodman, M.B., 2014. Mechanical control of the sense of touch by β-spectrin. Nat. Cell Biol. 16. doi:10.1038/ncb2915

Krieg, Michael, Dunn, A.R., Goodman, M.B., 2014. Mechanical control of the sense of touch by β-spectrin. Nat. Cell Biol. 16, 224–233.

Krieg, M., Stühmer, J., Cueva, J.G., Fetter, R., Spilker, K., Cremers, D., Shen, K., Dunn, A.R., Goodman, M.B., 2017. Genetic defects in β-spectrin and tau sensitize C. Elegans axons to movement-induced damage via torque-tension coupling. Elife 6. doi:10.7554/eLife.20172

McClanahan, P.D., Xu, J.H., Fang-Yen, C., 2017. Comparing Caenorhabditis elegans gentle and harsh touch response behavior using a multiplexed hydraulic microfluidic device. Integr.Biol. doi:10.1039/C7IB00120G

Nekimken, A.L., Fehlauer, H., Kim, A.A., Manosalvas-Kjono, S.N., Ladpli, P., Memon, F., Gopisetty, D., Sanchez, V., Goodman, M.B., Pruitt, B.L., Krieg, M., 2017a. Pneumatic stimulation of C. elegans mechanoreceptor neurons in a microfluidic trap 17, 1116–1127.

Nekimken, A.L., Mazzochette, E.A., Goodman, M.B., Pruitt, B.L., 2017b. Forces applied during classical touch assays for Caenorhabditis elegans. PLoS One 12. doi:10.1371/journal.pone.0178080

O’Hagan, R., 2005. Components of a Mechanotransduction Complex in C. elegans Touch Receptor Neurons: An in vivo Electrophysiology Study. Columbia University.

O’Hagan, R., Chalfie, M., Goodman, M.B., 2005. The MEC-4 DEG/ENaC channel of Caenorhabditis elegans touch receptor neurons transduces mechanical signals. Nat. Neurosci. 8, 43–50. doi:10.1038/nn1362

O’Toole, M., Lamoureux, P., Miller, K.E., 2015. Measurement of subcellular force generation in neurons. Biophys. J. 108, 1027–1037. doi:10.1016/j.bpj.2015.01.021

Petzold, B.C., Park, S.-J., Mazzochette, E.A., Goodman, M.B., Pruitt, B.L., 2013. MEMS-based force-clamp analysis of the role of body stiffness in C. elegans touch sensation. Integr. Biol. (Camb). 5, 853–864. doi:10.1039/c3ib20293c

Phillips, J.B., Smit, X., Zoysa, N. De, Afoke, A., Brown, R.A., 2004. Peripheral nerves in the rat exhibit localized heterogeneity of tensile properties during limb movement. J. Physiol. 557, 879–887. doi:10.1113/jphysiol.2004.061804

Ribeiro, A.J.S., Denisin, A.K., Wilson, R.E., Pruitt, B.L., 2016. For whom the cells pull: Hydrogel and micropost devices for measuring traction forces. Methods 94, 51–64. doi:10.1016/j.ymeth.2015.08.005

Sanzeni, A., Katta, S., Petzold, B., Pruitt, B.L., Goodman, M.B., Vergassola, M., 2019. Somatosensory neurons integrate the geometry of skin deformation and mechanotransduction channels to shape touch sensing. Elife 8, 1–44. doi:10.7554/eLife.43226

Stiernagle, T., 2006. Maintenance of C. elegans. WormBook. doi:10.1895/wormbook.1.101.1

Sure, G.R., Chatterjee, A., Mishra, N., Sabharwal, V., Devireddy, S., Awasthi, A., Mohan, S., Koushika, S.P., 2018. UNC-16/JIP3 and UNC-76/FEZ1 limit the density of mitochondria in C. elegans neurons by maintaining the balance of anterograde and retrograde mitochondrial transport. Sci. Rep. 8, 8938. doi:10.1038/s41598-018-27211-9

Suslak, T.J., Jarman, A.P., 2015. Stretching the imagination beyond muscle spindles - stretch - sensitive mechanisms in arthropods. J. Anat. 227, 237–242. doi:10.1111/joa.12329

Suzuki, H., Kerr, R., Bianchi, L., Frøkjær-Jensen, C., Slone, D., Xue, J., Gerstbrein, B., Driscoll, M., Schafer, W.R., 2003. In vivo imaging of C. elegans mechanosensory neurons demonstrates a specific role for the MEC-4 channel in the process of gentle touch sensation. Neuron 39, 1005–1017.

Tao, L., Porto, D., Li, Z., Fechner, S., Lee, S.A., Goodman, M.B., Xu, X.Z.S., Lu, H., Shen, K., 2019. Parallel Processing of Two Mechanosensory Modalities by a Single Neuron in C. elegans. Dev. Cell 51, 617–631.e3. doi:10.1016/j.devcel.2019.10.008

Vaadia, R.D., Li, W., Voleti, V., Singhania, A., Hillman, E.M.C., Grueber, W.B., 2019. Characterization of Proprioceptive System Dynamics in Behaving Drosophila Larvae Using High-Speed Volumetric Microscopy. Curr. Biol. 29, 935–944.e4. doi:10.1016/j.cub.2019.01.060

Vogel, B.E., Hedgecock, E.M., 2001. Hemicentin, a conserved extracellular member of the immunoglobulin superfamily, organizes epithelial and other cell attachments into oriented line-shaped junctions. Development 128, 883–894.

Zheng, Q., Ahlawat, S., Schaefer, A., Mahoney, T., Koushika, S.P., Nonet, M.L., 2014. The Vesicle Protein SAM-4 Regulates the Processivity of Synaptic Vesicle Transport. PLoS Genet. 10, e1004644. doi:10.1371/journal.pgen.1004644

