## Supplementary figures and images for "Touch-induced Mechanical Strain in Somatosensory Neurons is Independent of Extracellular Matrix Mutations in *C. elegans*"

### Supplemental Figures 1-3

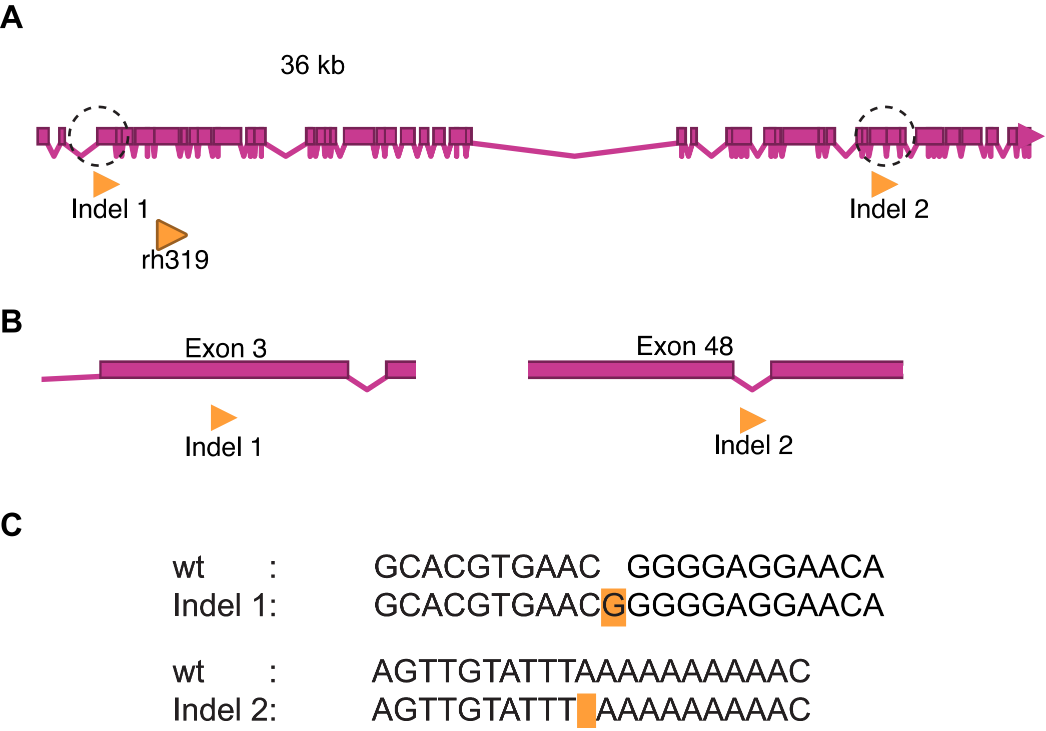

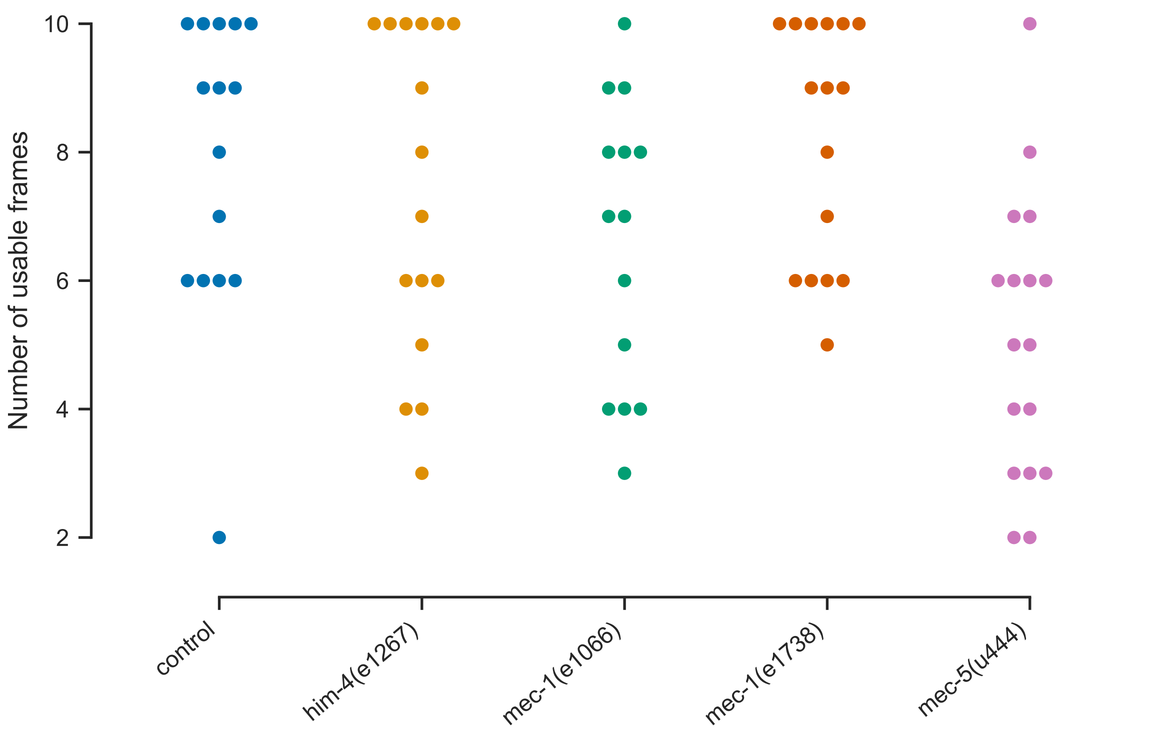

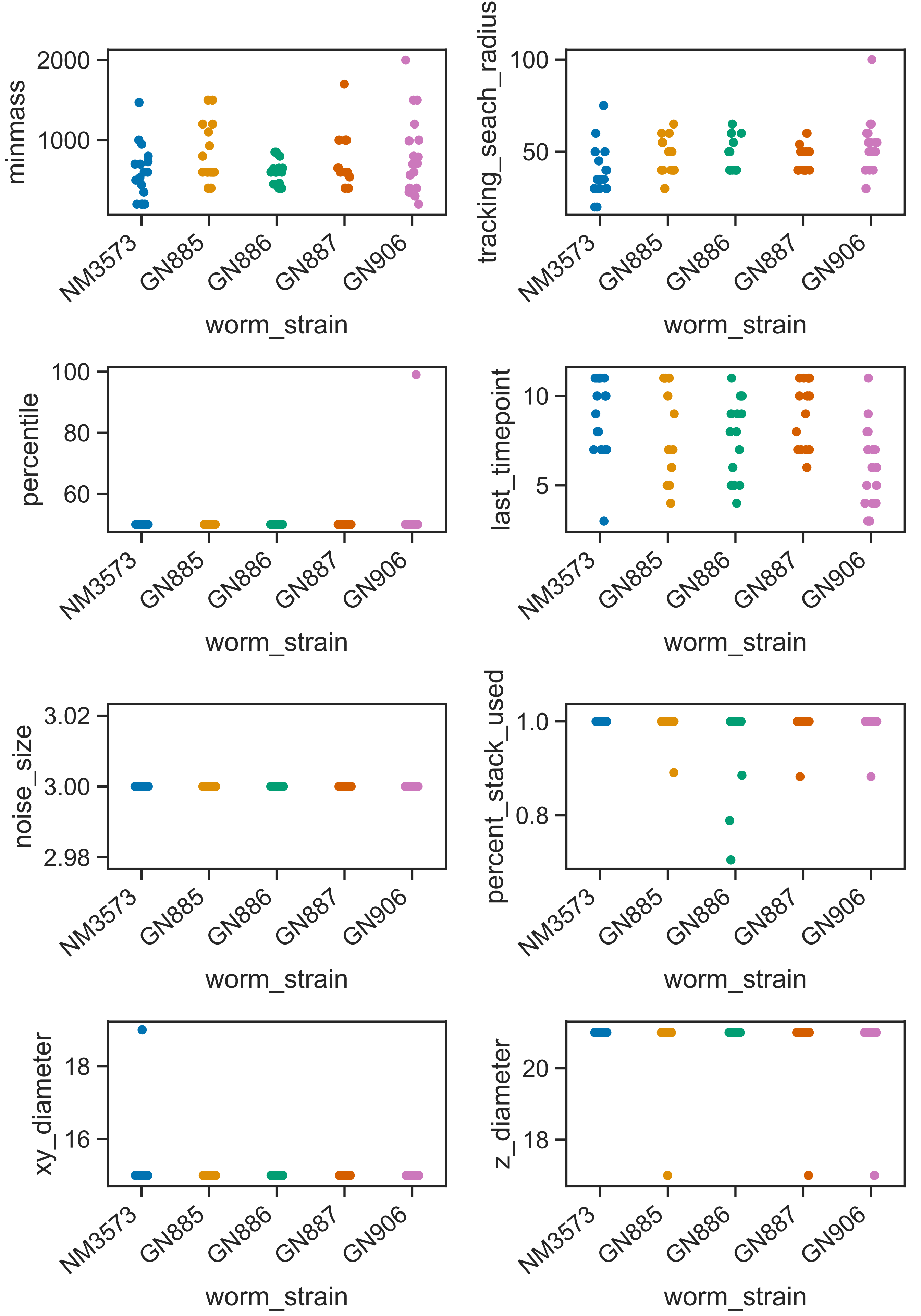
